# EOLA1, a novel mitochondria-localized protein critical for heart functions via regulating mitochondrial translation

**DOI:** 10.64898/2026.01.12.699056

**Authors:** Xiaoyan Shi, Yangyi Zhang, Nanbo Liu, Ruiqi Wang, Nan Zhang, Yanlan Cao, Dian Wang, Yuxuan Jin, Qingren Meng, Simin Fan, Jia Yao, Chih-hung Hsu, Shuoji Zhu, Ping Zhu, Yang Shi, Hao Chen

## Abstract

Mitochondria possess their own mRNA translation system, mediated by specialized mitoribosomes. Dysregulation of mitochondrial translation disrupts metabolic homeostasis and is linked to various pathologies, including metabolic syndromes, neurological disorders, and cardiac diseases. Using CRISPR/Cas9 screening, we identified EOLA1 (endothelial-overexpressed lipopolysaccharide-associated factor 1) as a novel mitochondrial matrix protein essential for metabolic homeostasis. Knockout of *Eola1* severely affects mitochondrial translation and consequently the oxidative phosphorylation in mammalian cells. Mechanistically, EOLA1 physically interacts with mitochondrial 12S ribosomal RNA (12S mt-rRNA) and the mitochondrial Tu translation elongation factor (TUFM), facilitating efficient translation elongation of mitochondrial-encoded mRNAs. Furthermore, genetic ablation of *Eola1* in mice induced severe cardiac dysfunction, demonstrating its indispensable role in maintaining heart function through mitochondrial translation control. Together, our findings establish EOLA1 as a key regulator of mitochondrial translation via its interactions with 12S mt-rRNA and TUFM, highlighting its importance in cellular metabolism and a potential role in cardiac disease pathogenesis.

## Introduction

Mitochondria are essential eukaryotic organelles of proteobacterial origin^1^, generating ATP via oxidative phosphorylation (OXPHOS), and playing critical roles in metabolic regulation, cellular signaling, and programmed cell death^2–4^. A notable difference between mitochondria and other intracellular organelles (except chloroplasts) is the presence of a completely independent mitochondrial genome and translation system in eukaryotic organisms^5–8^. In humans, mitochondrial DNA (mt-DNA) is a 16569 bp circular, double-stranded molecule encoding 37 essential genes^9^, including 13 protein-coding genes for core subunits of the electron transport chain (ETC), 22 tRNAs, and 2 rRNAs^5,10^ . While most mitochondrial proteins are nuclear-encoded and imported after cytosolic translation, the 13 mt-DNA-encoded proteins are vital for ATP production^6,7^. Notably, a significant proportion of the mitochondrial proteome is devoted to mt-DNA expression, highlighting the essential reliance on this compact yet indispensable genome for mitochondrial functions^6^.

Mitochondrial translation resembles prokaryotic systems but has also evolved unique features distinct from both bacteria and eukaryotic cytosol^6,11^. Most mitochondrial mRNAs (with the exception of two bicistronic transcripts) lack 5’ and 3’ untranslated regions (UTRs) and employ an ancient leaderless translation initiation mechanism^12^. Unlike nuclear DNA-encoded genes, mitochondrial gene expression exhibits spatiotemporal coupling of transcription, RNA maturation, translation, and RNA decay processes^13^. Notably, in contrast to the cytosolic translation machinery (requires at least 25 translation factors, some of which are multi-protein complexes), mitochondria possess a minimalistic set of translation factors^6,14,15^. These include initiation factors (mtIF2 and mtIF3), elongation factors (mtEF-G1, mtEF-G2, mtEF-Ts, and mtEF-Tu (also known as TUFM)), termination release factors (mtRF1a, mtRF1, mtRF-R, and ICT1), and additional translation regulators yet to be fully characterized^6^.

Mitochondrial translation is a fundamental process for cellular energy production, playing a vital role in maintaining cellular fitness and organismal health. Heart is an organ subjected to high energy requirements and depends heavily on mitochondria, as over 95% of ATP consumed by cardiomyocytes is derived from OXPHOS^16,17^. Consequently, mitochondrial translation dysfunction frequently manifests as cardiac pathologies, including cardiomyopathy and heart failure^12,18,19^. As a common and complex disease, cardiovascular diseases have become a significant global threat to human life and health^20,21^. The incidence and mortality rates have shown an overall upward trend in recent years, posing a serious public health challenge^20^. Therefore, identification of new molecules that regulate mitochondrial function and elucidating new mechanisms of metabolic disruption in the occurrence of heart diseases are of crucial importance for the prevention and treatment of cardiovascular diseases.

Endothelial-overexpressed lipopolysaccharide-associated factor 1 (EOLA1) is a 158-amino-acid protein harboring an ASCH domain (Extended Data Fig. 1a, b), which adopts a characteristic β-barrel fold surrounded by α-helices^22,23^. Initially identified as an inflammatory regulator, EOLA1 exerts its protective function through binding and stabilizing metallothionein-2A (MT2A)^24^, thereby suppressing pro-inflammatory mediators and dampening lipopolysaccharide (LPS)-induced signaling pathways^25,26^. Despite low sequence identity with other ASCH domains, EOLA1 retains a conserved “GxKxxExR” motif within its cavity structure, which harbors aromatic and polar residues predicted to accommodate nucleotides^22^. EOLA1 contains a positively charged cleft structurally similar to the YTH domain of m^6^A reader proteins and the ASCH domain of *Zymomonas mobilis* (zmASCH)^22,23^. The YTH domains recognize *N*^6^-adenosine RNA modifications^27^ while zmASCH domain exhibits single-stranded RNA binding and degradation activity *in vitro*^28,29^. Together, these conserved structural and functional features imply that EOLA1 may regulate gene expression through RNA binding during transcription coactivation, RNA processing, and regulation of translation.

In this study, we identified EOLA1 as a novel mitochondrial metabolism regulator through CRISPR/Cas9 screening. Bioinformatics and experimental validation revealed that EOLA1 is a mitochondrial matrix protein and interacts with the mitochondrial Tu translation elongation factor (TUFM) and 12S mitochondrial rRNA (mt-rRNA), forming a functional complex essential for mitochondrial mRNA translation elongation. Disruption of *Eola1* significantly impaired mitochondrial translation, reducing the synthesis of OXPHOS subunits and compromising ATP production. Remarkably, whole-body *Eola1* knockout in mice led to cardiac dysfunction, underscoring its physiological role in maintaining heart function via mitochondrial translation. Collectively, our study identifies EOLA1 as a novel mitochondria-localized protein serving as a critical regulator of mitochondrial protein synthesis and uncovers a novel mechanistic link between mitochondrial translational regulation and cardiovascular homeostasis, offering new insights into the pathogenesis of heart diseases.

## MATERIALS AND METHODS

### Cell culture

B16-F10 melanoma cells, HL-1 cardiac muscle cells and HEK293T cells were purchased from Wuhan Pricella Biotechnology. All cells were authenticated by STR profiling and testing for mycoplasma contamination was performed with a MycAway one-step mycoplasma detection kit (Yeasen). Unless otherwise noted, all cells were cultured in Dulbecco’s Modified Eagle Medium (DMEM, Gibco) containing 10% fetal bovine serum (FBS, BI) and 1% penicillin-streptomycin (Thermo Fisher Scientific) and maintained in a 37°C humidified incubator with 5% CO_2_.

### Plasmid construction

To express double-tagged FLAG–HA human EOLA1 protein (EOLA1–FH) into HEK293T cells, EOLA1 cDNA was cloned into the retroviral vector pHAGE. Truncated ORF DNA of EOLA1 (amino acids position 30 ∼ 158) was inserted into the retroviral vector pHAGE to express EOLA1 protein that lacks mitochondria-targeting signals (MTS). To generate *Eola1* knockout cell lines, *Eola1*-specific 20nt small guide RNA (sgRNA) sequences designed to target the second exon of *Eola1* were cloned into pL-CRISPR.EFS.GFP vector.

sgRNAs targeting mouse *Eola1* locus:

sgRNA: ACAGATTCAGACCTTACTACAGG

### Generation of EOLA1 knockout and overexpression stable cell lines

To generate *Eola1* knockout cells, 2 μg of sgRNA plasmid was transfected to B16-F10 and HL-1 cells using Lipofectamine 3000 (Invitrogen). After 2 days’ culture in DMEM medium, cells were screened via fluorescence and isolated into monoclons by flow cytometry. After 3 weeks’ culture, colonies were screened out and confirmed by Western blotting with EOLA1 antibody and sanger sequencing (cloned into T-vector)). To generate EOLA1 overexpression stable cell lines, the pHAGE empty vector, EOLA1–FH, or truncated EOLA1–FH plasmid was co-transfected with pMD2.G and PsPAX2 packaging plasmids into HEK293T cells, respectively. After 7 days’ selection with 2 μg/mL puromycin, cells were confirmed by Western blotting with EOLA1 antibody.

### CRISPR/Cas9 library screening

To generate Cas9-stable B16-F10 melanoma cells, we introduced a lentiviral vector encoding both Cas9 and a fluorescent reporter protein, followed by fluorescence-based cell sorting. For individual gene knockdowns, we stably transfected sgRNAs into these Cas9-expressing cells and selected with puromycin (2 μg/mL, 7 days). To perform phenotypic screening, sgRNA-transduced cells were cultured for 48 hours in either galactose medium (10 mM galactose, 1 mM pyruvate, 2 mM glutamine, 10% FBS, 1% penicillin-streptomycin) or glucose medium (10 mM Glucose, 1 mM pyruvate, 2 mM glutamine, 10% FBS, 1% penicillin-streptomycin). Subsequently, dead cells from galactose and living cells from glucose conditions were collected for genomic DNA (gDNA) extraction. sgRNA fragments were amplified from 5 μg gDNA using 2× PCR Master Mix, and the resulting libraries were subjected to high-throughput sequencing (Berry Genomics) for CRISPR screen analysis.

The CRISPR/Cas9 screening data analysis was performed by MAGeCK (v0.5.9.5) as previously described^30^. MAGeCK RRA method was used to evaluate the abundance of sgRNAs and identify gene hits.

### Apoptosis and cell proliferation assay

For apoptosis analysis, a 6-well plate of cells cultured in glucose or galactose medium was harvested by treatment with trypsin without EDTA (Gibco). The staining was performed by the use of an Annexin V-FITC/PI Apoptosis Detection Kit (Vazyme), and the apoptotic cells were detected by flow cytometry analysis (BD FACSCanto). For cell proliferation assay, 2 × 10^5^ cells were plated on one well of a 24-well plate in glucose-based or galactose-based medium on day 0 and counted from day one until day 3 with cell counters (Thermo Fisher Scientific).

### Mitochondrial isolation and Protease K protection assay

Cells were collected and resuspended with 5 ml MTiso-buffer (3 mM HEPES-KOH pH 7.4, 210 mM mannitol, 70 mM sucrose, 0.2 mM EGTA, Complete Mini EDTA-free) and then transfer the suspension into Dounce homogenizer for 50 stokes. The supernatant was centrifuged at 800 g for 10 min at 4 °C to remove nuclei and unbroken cells and then centrifuged at 15000 g for 10 min at 4 °C to collect the mitochondrial pellet. Protease K protection assay was performed as previously described^31^. In brief, 0.5 mg mitochondria from HEK293T cells were resuspended within the three following buffers: (1) Iso-osmotic (10 mM MOPS-KOH pH 7.2, 250 mM sucrose, 1 mM EDTA), (2) Hypo-osmotic (10 mM MOPS-KOH pH 7.2, 1 mM EDTA) and (3) Membrane perforated (10 mM Tris-HCl pH 7.5, 1mM EDTA, 1% Triton X-100). Proteinase K (20 mg/mL) was then added in. After 30 min of incubation on ice, the reaction was stopped by adding 20 mM PMSF. All three reaction sets were confirmed by Western blotting with corresponding antibodies.

### Fluorescence microscopy

To verify the mitochondrial localization of EOLA1 protein, HEK293T stably expressing EOLA1-FH or Truncated EOLA1-FH were used for immunofluorescence. Cells seeded on coverslips early were fixed by incubating with 4% (w/v) paraformaldehyde for 10 min at room temperature. Subsequently, cells were permeabilized by incubating with 0.5% Triton X-100 for 15 min and blocked by incubating for 1 hour in 1% (w/v) BSA. Cells were then incubated with the corresponding antibodies overnight at 4 °C and washed three times before the secondary antibody incubation. Fluorescence microscopy images were taken by using an inverted LSM980 Laser Scanning Confocal Microscope (LSM 980 with Airyscan). The corresponding antibodies included anti-FLAG antibodies (1:1000, Sigma, F1804), anti-HA antibodies (1:1000, Abcam, ab314237), anti-HSP60 antibodies (1:3000, Proteintech, 15282-1-AP), anti-TFAM antibodies (1:1000, Proteintech, 22586-1-AP), anti-mouse Alexa Fluor 488 (1:2000, Invitrogen, A-10680), and anti-rabbit Alexa Fluor 555 (1:2000, CST, 4413).

### Complex purification

To purify EOLA1 complex, four 15-cm plates of 293T cells stably expressing FLAG-HA (FH) tag alone (mock) or 293 cells stably expressing EOLA1–FH were collected and then lysed on ice with 500 µL IP buffer (10 mM Tris–HCl pH 7.5, 100 mM KCl, 5 mM MgCl_2_, 1 mM DTT, 0.25% NP40% and protease inhibitor) for 10 min. Protein extracts were collected via centrifugation at 12000 g for 10 min at 4 °C and then incubated with anti-FLAG beads (Invitrogen) overnight for immunoprecipitation. Beads-protein mixture was washed five times with IP buffer before elution, which then was performed by competition with FLAG peptides. Eluted proteins were tested by silver and subsequently subjected to high-performance liquid chromatography-mass spectrometry (HPLC-MS) to analyze EOLA1-associated factors.

### Western blotting

For western blotting, a six-well plate of cells at approximately 80-90% confluency was lysed with 100 μl RIPA buffer on ice. Protein extracts were separated by 4-20% SDS-PAGE, and the protein was then transferred to a 0.4 μm polyvinylidene fluoride (PVDF, Merck Millipore) membrane. After blocking with 5% milk in TBST, the membranes were then incubated with the corresponding antibodies overnight at 4°C. Membranes were incubated with secondary HRP-conjugated antibody (Proteintech) at room temperature for 1 hour following three times washes with TBST and were washed three times again before being detected by the phosphoimager.

The corresponding antibodies included anti-EOLA1 antibody (1:5000, CUSABIO, CSB-PA837446LA01HU), anti-HSP60 antibody (1:3000, Proteintech, 15282-1-AP), anti-Tom20 antibody (1:1000, ABclonal, A19403), anti-α-Tubulin antibody (1:3000, Proteintech, 80762-1-RR), anti-HistoneH1.2 antibody (1:1000, Proteintech, 19649-1-AP), anti-TUFM antibody (1:1000, SAB, 34669), anti-FLAG antibody (1:1000, Sigma, F1804), anti-MRPS15 antibody (1:1000, Proteintech, 17006-1-AP), anti-MRPL11 antibody (1:1000, Proteintech, 15543-1-AP), anti-ATP6 antibody (1:500, SAB, 31464), Anti-COX1 antibody (1:500, ABclonal, A23123), anti-MT-ND5 antibody (1:500, Proteintech, 55410-1-AP), anti-β-actin antibody (SAB, 52901), and anti-GAPDH antibody (1:5000, Proteintech, 10494-1-AP).

### Seahorse

The oxygen consumption rate (OCR) measurements were performed on WT and EOLA1 KO cells using an XF96 extracellular flux analyzer (Agilent), where 40000 cells per well were seeded, cultured overnight in 5% CO_2_ at 37°C, washed three times with assay medium, equilibrated for 1 hour in a CO_2_-free incubator, and assessed following sequential injection of oligomycin (1 μM), FCCP (1.5 μM), and rotenone/antimycin A (0.5 μM).

### Mitochondrial polysome profiling

Two 15-cm plates of HL-1 cells prepared for each sample (WT and EOLA1 KO) were treated with 100 µg/mL cycloheximide and incubated for 8 min at 37 °C. Cells were lysed on ice for 10min in 400 µL lysis buffer (20 mM Tris–HCl pH 7.5, 150 mM NaCl, 5 mM MgCl_2_, 1 mM DTT, 100 μg/mL Cycloheximide, 1% Triton X-100). The supernatant was collected via centrifugation at 12000 g for 10 min at 4 °C, and its’ absorbance at 260 nm was measured. The ribosomes with equal A260 in each group were loaded onto a sucrose gradient (10–30%) buffer prepared using a gradient station and centrifuged in an SW41Ti rotor (Beckman) at 4°C for 4 hours at 35000 rpm. Polysome profiles were then generated and analyzed by Gradient Station. A total of 16 fractions were collected from the top and RNA samples were extracted by TRIzol reagent (Invitrogen) and reverse transcribed followed by Quantitative real-time PCR (RT-PCR) detection. Primer sequences are provided in Supplementary Table 1.

### Quantitative real-time PCR (RT-PCR)

Total RNA was extracted using TRIzol reagent (Invitrogen). Reverse transcription was performed with 1 mg of total RNA using the HiScript III RT SuperMix for qPCR (Vazyme). RT-PCR was conducted using ChamQ SYBR qPCR Master Mix (Vazyme) on a QuantStudio 3 Real-Time PCR System (Applied Biosystems). Primer sequences are provided in Supplementary Table 1.

### UV-RNA Immunoprecipitation (UV-RIP) sequence

UV-RNA-seq was performed as previously described^32^. In brief, a 10-cm plate of 293 cells were washed with ice-cold PBS containing protease inhibitors, UV-crosslinked at 400 mJ/cm², and collected by scraping. The pellet was lysed sequentially in low-salt lysis buffer (50 mM Hepes-KOH pH 7.5, 10 mM NaCl, 1 mM EDTA, 10% glycerol, 0.2% NP-40, 1% Triton X-100) and high-salt lysis buffer (600 mM NaCl, 10 mM Tris-HCl pH 8.0, 1 mM EDTA, 0.5 mM EGTA, 1% Triton X-100, 0.1% DOC). Lysates were incubated with EOLA1 antibody in IP buffer (10 mM Tris-HCl pH 8.0, 1 mM EDTA, 0.5 mM EGTA, 1% Triton X-100, 0.1% DOC) on a rotating platform overnight at 4 °C. Beads were washed sequentially with low-salt wash buffer (150 mM NaCl, 0.1% SDS, 1% Triton X-100, 2 mM EDTA, 20 mM Tris-HCl pH 8.0), high-salt wash buffer (500 mM NaCl, 0.1% SDS, 1% Triton X-100, 2 mM EDTA, 20 mM Tris-HCl pH 8.0), and LiCl wash buffer (0.25 M LiCl, 1% IGEPAL CA630, 1% sodium deoxycholate, 1 mM EDTA, 10 mM Tris pH 8.0), followed by TE buffer. RNA–protein complexes were eluted in elution buffer (10 mM Tris–HCl pH 8.0, 1 mM EDTA, 1% SDS) and RNA extracted by TRIzol reagent (Invitrogen). Purified RNA underwent PNK treatment, followed by library preparation with the VAHTS^TM^ Small RNA Library Prep Kit for Illumina^®^ (Vazyme).

### RIP-qPCR

The RNA immunoprecipitation experiment was conducted according to manufacturer’s instructions with minor modifications. Briefly, 1×10^7^ cells in a 150-mm dish were subjected to RIP-qPCR. Cell pellets were lysed in RIP buffer (25 mM Tris–HCl pH 7.5, 150 mM KCl, 2 mM EDTA, 0.5% NP40, 0.5 mM DTT, 1 X protease inhibitor cocktail). The mRNAs were then pulled down by antibody following the immunoprecipitation protocol. The co-precipitated RNA was recovered from the beads using a RNeasy mini kit (Qiagen Inc). Both the input and co-immunoprecipitated RNA were recovered using an RNA Clean & Concentrator-5 kit (Zymo) and subjected to RT-qPCR analyses. Primer sequences are provided in Supplementary Table 1.

### Mito-Click-iT assay

Cells were grown to approximately 80–90% confluency, followed by emetine (50 μg/mL) treatment to inhibit cytoplasmic protein synthesis and labeling of mitochondrial proteins with L-AHA (Invitrogen, C10102, 25–50 μM) for 1–4 hours. Mitochondria were isolated using the mitochondrial isolation method described above. The isolated mitochondria were resuspended in 0.2 mL of Click lysis buffer (1% SDS, 50 mM Tris-HCl, pH 8.0), and protein concentration was measured using the Pierce™ BCA Protein Assay Kit (Thermo Fisher Scientific). The click reaction was performed using the Click-iT® Cell Reaction Buffer Kit (Thermo Fisher Scientific) and incubated for 1–1.5 hours. Proteins were denatured with 20 µL of 6X protein loading buffer at 95°C for 5 minutes. Western blotting detection is performed using streptavidin-HRP for biotinylated nascent proteins.

### Animals

Whole-body *Eola1* knockout (Eola1^-/-^) mice were generated on a C57BL/6N background. Both male and female mice at five weeks of age were included for phenotypic analysis. All mice were bred and maintained under specific-pathogen-free (SPF) conditions at the Animal Experiment Center of Southern University of Science and Technology (SUSTech). Animal experiments were conducted in accordance with protocols approved by the SUSTech Institutional Animal Care and Use Committee (IACUC).

### Echocardiography

To evaluate the major structures and functions of the mouse heart in vivo, we performed standardized transthoracic echocardiographic examinations. Mice were anesthetized with 1.5% isoflurane and examined in a blinded manner using a 40 MHz probe (MS400) of the Vevo2100 echocardiographic imaging system (VisualSonics, Toronto, Ontario, Canada). Both motion mode (M-mode) and brightness mode (B-mode) were used to record ultrasound imaging videos of myocardial tissue contraction and relaxation. The long axis, short axis, and area of the left ventricle were identified and calculated on the B-mode images at the end of diastole. The anterior and posterior walls of the left ventricle were identified on the M-mode images, and ejection parameters and wall thickness based on four consecutive cardiac cycles were automatically calculated. All cardiac physiological parameter measurements were performed offline using VevoLab analysis software (FUJIFILM VisualSonics, Toronto, Ontario, Canada).

### Statistical analysis

No sample size calculations and blinding were performed. There was no method of randomization. No samples or animals were excluded from analysis. Prism 9 software (GraphPad) was used to analyses data from biological experiments by performing two-tailed unpaired Student’s t-tests. One-way analysis of variance (ANOVA) with Tukey’s test was used to compare multiple groups and two-way ANOVA was used to compare cell proliferation analysis. *P* values \**P*<0.05, \*\**P*<0.01, \*\*\**P*<0.001, \*\*\*\**P*<0.0001 were considered statistically significant.

## RESULTS

### A CRISPR/Cas9 screen identified EOLA1 as a novel regulator of mitochondrial activity

To systematically identify novel regulators of mitochondrial function, we performed an unbiased positive CRISPR/Cas9 screening focused on 934 poorly characterized protein-coding genes using a 10076 unique single-guide RNAs (sgRNAs) library in B16-F10, a murine cell model of melanoma. It was reported that mitochondrial activity can be rapidly assessed by monitoring cell apoptosis and proliferation in the medium that contains galactose instead of glucose as the primary carbon source^33^. As the catalytic reaction of galactose to glucose-1-phosphate is very slow and ATP production by glycolysis is severely restricted, OXPHOS becomes the sole manner to generate ATP^34^. Thus, cells with mitochondrial dysfunction cannot proliferate or even survive in galactose medium. To identify key mitochondrial metabolic regulatory genes by exploiting this metabolic vulnerability, B16-F10 melanoma library cells were cultured under two conditions: glucose medium or galactose medium (galactose replacing glucose as the primary carbon source) for 48 hours. Control cells from glucose medium or dead cells from galactose medium were collected to carry out next-generation sequencing for evaluating the abundance of sgRNAs (Fig. 1a). Then, MAGeCK^30^ algorithm were employed to identify essential genes in mitochondrial activity. As shown in Fig. 1b, several of the top hit genes have been demonstrated to be directly or indirectly involved in OXPHOS-linked energy metabolism, e.g., *Sp1*^35^, *Stat1*^36^, *Samhd1*^37^ and *Stim1*^38^, which indicates that our screening is of high quality, validity and reliability. Among the rest top candidate genes, *Eola1* emerges as a compelling target due to its ranking score while with barely characterized biological functions. This knowledge gap, along with its pronounced expression in our screening data, highlights *Eola1* as a high-priority candidate for functional characterization, particularly its potential role in mitochondrial regulation (Fig. 1b, c).

**Fig. 1:**
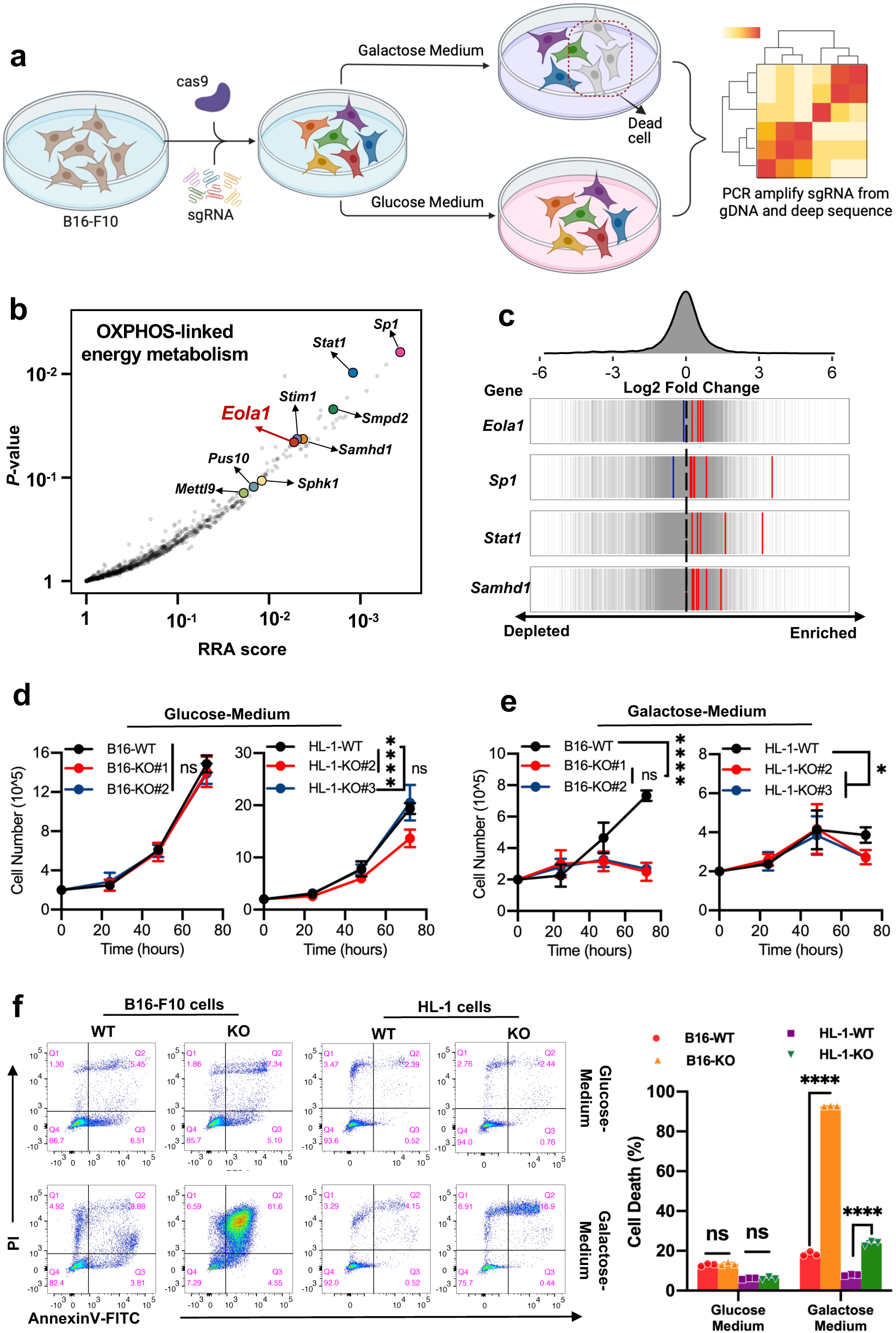
CRISPR/Cas9 screening identifies essential genes in the regulation of mitochondrial function. **a**, Schematic diagram of screening strategy. Cell populations with sgRNA library were cultured in glucose medium or galactose medium (galactose was used to replace glucose) for 48 hours. Dead cells from galactose medium or living cells from glucose medium were collected to carry out next-generation sequencing for evaluating the abundance of sgRNAs. **b**, Genes essential for mitochondrial oxidative phosphorylation (OXPHOS) -linked energy metabolism were identified in the screening. A rank-ordered plot shows the RRA (Robust Rank Aggregation) scores and corresponding *p*-values from the CRISPR screening results. **c**, Fold change (FC) distribution of sgRNAs targeting *Eola1, Sp1, Stat1,* and *Samhd1* as indicated in red (upregulated) and blue (downregulated) lines, overlaid on gray gradient depicting the overall distribution. **d**, **e**, Growth curves of wild-type (WT) and EOLA1 KO cells cultured in the medium of glucose (**d**) or galactose (**e**). **f**, *Eola1* depletion dramatically induced cell apoptosis when cancer cells cultured in galactose medium. Representative flow cytometry images (left) and quantification analysis (right) of the apoptotic B16-F10 melanoma cells and HL-1 cells cultured in glucose medium or galactose medium for 48 hours. FITC: fluorescein isothiocyanate; PI: propidium iodide. **Note: d**, **e**, One-way ANOVA with Tukey’s test. **f**, two-tailed unpaired Student’s t-tests. \**P*<0.05, \*\*\*\**P*<0.0001, ns: not significant. **d**, **e**, **f**, Data were presented as mean ± SD (n=3).

To validate the screening results, we generated *Eola1* knockout B16-F10 melanoma cells using CRISPR/Cas9 (Extended Data Fig. 1c-e) and compared their growth and apoptotic status to wild-type (WT) cells in galactose- or glucose-based medium over 48 hours. Consistent with high-throughput screening results, *Eola1* knockout cells exhibited significantly suppressed proliferation and increased apoptosis when cultured in galactose medium. In contrast, no notable changes in cell growth or apoptosis were observed when cultured in glucose medium (Fig. 1d-f). To further verify the screening results, we chose another distinct cell line, HL-1 cardiac muscle cell, to detect the essentiality of *Eola1* for mitochondria. Similar to the observations in B16-F10 cell line, *Eola1* depletion in HL-1 cells led to a marked increase in apoptosis and suppressed proliferation when cultured in galactose medium, with no substantial impact to the cell growth in glucose medium (Fig. 1d-f). These results collectively suggest that EOLA1 plays a critical role in maintaining mitochondrial function.

### EOLA1 exclusively localizes in mitochondria and is required for mitochondrial activity

Mitochondria are key organelles containing over 1000 proteins of dual genetic origin^3,39^. Given the essentiality of EOLA1 for mitochondrial activity, we investigated whether it is a mitochondrial protein but not a nuclear protein as previously reported^26^. First, we searched MitoCarta 3.0, a comprehensive database of mammalian mitochondrial proteins, but found no record of EOLA1. We then employed computational method (TargetP 2.0) to predict mitochondrially targeted proteins that might be missed in MitoCarta 3.0 database. In total, we identified 514 evolutionarily conserved mitochondrial proteins, along with 144 human-specific and 129 mouse-specific candidates (Fig. 2a). Cross-referencing the 514 shared proteins with the MitoCarta 3.0 database (1136 proteins) revealed that most of them are overlapped with established mitochondrial proteins, yet 27 predicted proteins were absent in MitoCarta 3.0 (Fig. 2a). Importantly, while these 27 proteins were unannotated in MitoCarta, literature evidences support the mitochondrial localization of several candidates, including MP31^40^, GAL3A^41,42^ and SARM1^43,44^. Interestingly, EOLA1 was also among these 27 high-confidence while unannotated mitochondrial candidates. Given that most of nuclear-encoded mitochondrial proteins are typically imported via an N-terminal mitochondrial-targeting sequence (MTS)^3,45^, we further analyzed the sequence of EOLA1 across diverse species and found a well-conserved MTS at its N-terminus (Extended Data Fig. 2a). These features strongly support EOLA1’s potential function as a mitochondria-localized protein. However, a previous study reported that it is localized in both the cytoplasm and nucleus of ECV304 cells^26^. To experimentally clarify the intracellular distribution of the EOLA1 protein, we firstly carried out the immunofluorescence experiment and observed an exclusive mitochondrial localization of EOLA1, which is well overlapped with the staining pattern of known mitochondrial proteins (Fig. 2b). Furthermore, our analysis unveiled that the N-terminus-truncated EOLA1 apparently lost the capacity to enter mitochondria (Extended Data Fig. 2b), which suggests that its predicted MTS is a functional element for mitochondrial importing of EOLA1. Consistently, subcellular fractionation combined with Western blotting confirmed the mitochondrial localization of the endogenous EOLA1 in both *Homo sapiens* and *Mus musculus* (Fig. 2c).

**Fig. 2:**
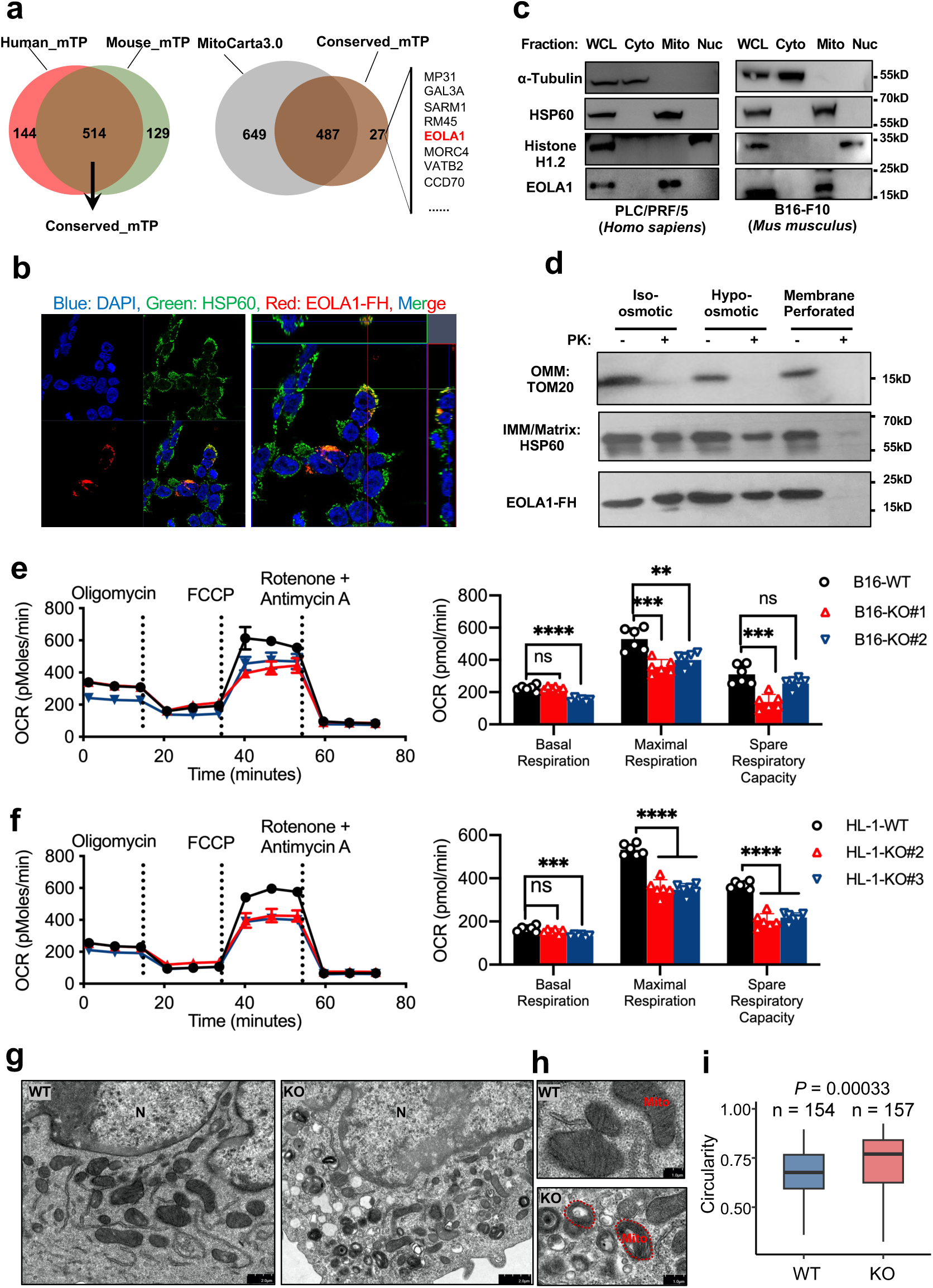
EOLA1 is a novel mitochondrial protein and required for mitochondrial activity. **a,** Mitochondrial proteins predicted by TargetP 2.0 in human and mouse (left), and their overlap with MitoCarta 3.0 annotations (right). **b,** Fluorescence analysis of the subcellular location of human EOLA1 protein (red), and HSP60 (green) was used as a marker for mitochondria. The exogenously expressed EOLA1 was tagged with a C-terminal FLAG and HA tandem epitope. **c,** Western blotting analysis of EOLA1 in the subcellular fractions of PLC/PRF/5 (*Homo sapiens*, left) and B16-F10 melanoma cells (*Mus musculus*, right). Tubulin was used as a cytoplasmic marker, Histone H1.2 as a nuclear marker, and HSP60 as a mitochondrial marker. **d,** Determination of EOLA1 sub-mitochondrial localization by Proteinase K digestion assay using purified mitochondria from HEK293T cells. PK, Protease K; IMM, inner mitochondrial membrane; OMM, outer mitochondrial membrane. **e, f,** Seahorse analysis of WT and *Eola1* KO in B16-F10 melanoma cells (**e**) and HL-1 cells (**f**). Quantification of the basal respiration, maximal respiration, and spare respiratory capacity was performed for the indicated cells. **g-i,** Electron microscopy images (**g**) of WT cells (left) and *Eola1* knockout cells (right). Higher-magnification views are presented in panel **h**. The relative circularity ratio of mitochondria was quantified (**i**). Mito: Mitochondria, N: nucleus. **Note: e**, **f**, One-way ANOVA with Tukey’s test. **i**, two-tailed unpaired Student’s t-tests. \*\**P*<0.01, \*\*\**P*<0.001, \*\*\*\**P*<0.0001, ns: not significant. **e**, **f**, Data were presented as mean ± SD (n=6).

The mitochondrion, a double-membrane-bound organelle, comprises three distinct compartments: the outer membrane, inner membrane, and mitochondrial matrix^3^. To determine the precise sub-mitochondrial localization of EOLA1, we performed a Proteinase K protection assay in combination with Western blotting. The results revealed that EOLA1 was barely digested by Proteinase K unless the outer and inner mitochondrial membranes are destroyed, suggesting its localization within the mitochondrial matrix (Fig. 2d). Since EOLA1 is a mitochondrial protein and mitochondria play a central role in generating cellular ATP via OXPHOS, we next investigated whether EOLA1 regulates mitochondrial respiration. Using the Seahorse XF Analyzer, we measured the oxygen consumption rate (OCR) in WT and *Eola1* knockout cells to quantitatively assess mitochondrial respiratory function. Notably, *Eola1* KO cells exhibited a significantly reduced OCR compared to WT cells (Fig. 2e, f), indicating that EOLA1 is essential for efficient OXPHOS activity. Moreover, mitochondria sustain cellular functions by dynamically modulating their morphology and architecture^46^. Consistent with this adaptive plasticity, electron microscopy revealed that mitochondria in WT cells maintained the typical ovoid shape, with intact double membranes and well-organized cristae. In contrast, *Eola1* KO mitochondria exhibited rounding, a significant reduction in length, and diminished cristae density, indicating deteriorated mitochondrial function^47,48^ (Fig. 2g-i). Collectively, our findings here provide evidence that EOLA1 is a *bona fide* mitochondrial protein mainly residing in the internal matrix, and plays a critical role in maintaining mitochondrial function.

### EOLA1 interacts with TUFM and 12S mt-rRNA

Why is EOLA1 so critical for mitochondria? To address this question, we combined a series of molecular and biochemical assays to delineate the mechanistic link between *Eola1* deficiency and mitochondrial impairment. Firstly, we generated stable 293T cells expressing EOLA1 tagged with FLAG-HA epitopes at its C-terminus and utilized the affinity purification coupled with liquid chromatography-tandem mass spectrometry (LC-MS/MS) to identify EOLA1-interacting partners. Remarkably, over 35% of its interacting partners were mitochondrial proteins (Extended Data Fig. 3a), suggesting a predominant role in mitochondrial processes. Among these, we specifically identified TUFM (Fig. 3a), a key regulator of mitochondrial protein translation^6^. Importantly, this interaction was readily confirmed by Western blotting using anti-TUFM antibodies (Fig. 3b). Given that EOLA1 harbors an ASCH domain (Extended Data Fig. 1a, b), which is predicted to bind nucleotides^22,23^, we hypothesized that it might also interact with mitochondrial RNAs to modulate translation. To test this hypothesis, we performed UV-RIP-seq^32^, which revealed robust enrichment of 12S mitochondrial rRNA (mt-rRNA) in EOLA1 pull-downs (Extended Data Fig. 3b). Notably, 12S mt-rRNA serves as the essential RNA component of the mitochondrial ribosome^6^, further supporting EOLA1’s potential role in mitochondrial translation regulation. In line with UV-RIP-seq results, exogenous RIP-qPCR confirmed EOLA1’s specific binding to 12S mt-rRNA under both UV-crosslinked and non-crosslinked conditions (Fig. 3c, d). Furthermore, endogenous RIP-qPCR also independently validated this interaction, demonstrating its physiological occurrence (Fig. 3e). Taken together, these data suggested that EOLA1 may play a role in regulating mitochondrial translation via interacting with TUFM and 12S mt-rRNA.

**Fig. 3:**
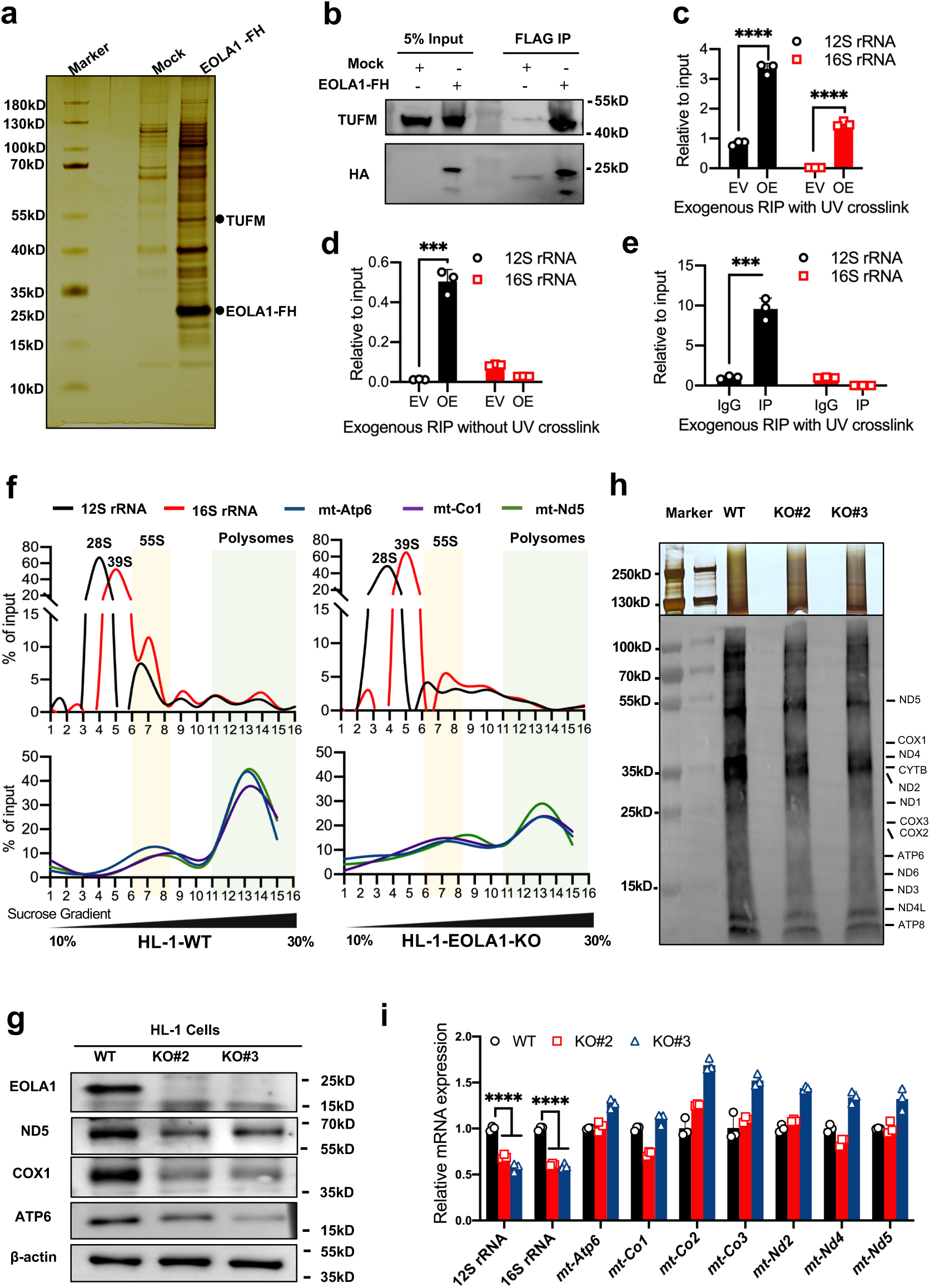
EOLA1 regulates mitochondrial mRNA translation by interacting with TUFM and 12S mt-rRNA. **a, b,** Identification of EOLA1 interacting partners. EOLA1 complex was purified from EOLA1–FH stable cells using immunoaffinity purification. Eluted proteins were separated by SDS-PAGE and analyzed by silver staining (**a**). Western blotting analysis verified the mitochondrial Tu translation elongation factor (TUFM) as an EOLA1-binding partner (**b**). **c-e,** RNA immunoprecipitation (RIP) followed by qPCR demonstrates specific binding between EOLA1 and 12S mt-rRNA. Exogenous RIP-qPCR with (**c**) or without (**d**) UV crosslinking confirms direct RNA-protein interactions. Endogenous RIP-qPCR further validates this physiological association (**e**). The identification primers are provided in supplementary Table 1. **f, g,** EOLA1 affects mitochondrial mRNA translation. Sucrose gradient fractionation profiles of mitochondrial ribosomal subunits assessed by RT-PCR of 12S rRNA (mt-SSU) and 16S rRNA (mt-LSU) (**f**, top). Distribution of mitochondrial-encoded mRNAs (ATP6, ND5, and COX1) measured by RT-PCR (**f**, bottom), along with their corresponding protein levels measured by Western blotting. β-actin served as a loading control (**g**). The identification primers are provided in supplementary Table 1. **h,** Mitochondrial translation activity was assessed using the Mito-Click-iT assay. **i,** Mitochondrial-encoded mRNAs (mt-mRNAs) and ribosomal RNA levels (12S and 16S mt-rRNA) were quantified in control and EOLA1-knockout cells by RT-qPCR. The identification primers are provided in supplementary Table 1. **Note: c**, **d**, **e**, two-tailed unpaired Student’s t-tests. **i**, One-way ANOVA with Tukey’s test. \*\*\**P*<0.001, \*\*\*\**P*<0.0001. **c**, **d**, **e**, **i**, Data were presented as mean ± SD (n=3).

### Depletion of EOLA1 affects protein synthesis in mitochondria

To elucidate the mechanistic link between EOLA1-TUFM/mt-rRNA interactions and mitochondrial translation regulation, we performed systematic analysis of mitoribosome assembly and translation efficiency. First, we carried out mitochondrial polysome profiling combined with RT-qPCR in HL-1 cardiac muscle cell to assess ribosomal subunit distribution (Fig. 3f and Extended Data Fig. 3c). Quantification of 12S and 16S mt-rRNA across a 10–30% sucrose gradient was used to monitor the dynamic distribution of mitochondrial ribosomal small subunits (mt-SSUs) (the first peak of 12S rRNA, fractions 4), large subunits (mt-LSUs) (the first peak of 16S rRNA, fractions 5), monosomes (the co-fractionated peak, fractions 7), and polysomes (co-fractionated peaks, fractions 11-15) between WT and EOLA1 KO cells. Compared to WT cells, KO cells showed significantly reduced ratios of mt-SSUs and co-fractionation of 12S and 16S mt-rRNAs (Fig. 3f), thus indicating a major defect in mt-mRNA translation. There are thirteen mt-DNA-encoded proteins synthesized by mitochondrial ribosomes. Therefore, we also analyzed the distribution of three representative mRNAs (ATP6, ND5, and COX1) encoded by the mitochondrial genome and observed a striking reduction in the fractions of mature ribosomes in *Eola1*-depleted cells compared with WT cells, which indicated that the active translation of these mRNAs was severely repressed (Fig. 3f). Consistently, Western blotting analysis confirmed substantially reduced protein levels of ATP6, ND5, and COX1 in KO cells, suggesting that EOLA1 is critical for mt-mRNA translation (Fig. 3g). To directly measure translation output, we employed Mito-Click-iT assay, which also confirmed significantly diminished mitochondrial protein synthesis in KO cells (Fig. 3h). Interestingly, significant reduction of 12S and 16S mt-rRNA was observed in EOLA1 KO cells (but we didn’t detect any obvious alterations in the mt-mRNA levels), which may be due to compromised structural stability of mitoribosome (Fig. 3i). Collectively, these data demonstrate that EOLA1 critically regulates mitochondrial translation by forming a functional complex with TUFM and 12S mt-rRNA.

### *Eola1* deletion results in dilated cardiomyopathy

To investigate EOLA1’s biological functions, we generated whole-body *Eola1* knockout mice on a C57BL/6 background (Fig. 4a and Extended Data Fig. 4a, b) and found that the knockout mice displayed no overt developmental abnormalities, including normal reproduction, gross morphology, and body weight (Extended Data Fig. 4c, d). Given the heart’s exceptionally high energy demands, with cardiomyocytes obtaining over 95% of their ATP from mitochondrial OXPHOS^6,16,17^, we hypothesized that EOLA1 may play a crucial role in cardiac metabolism. Additionally, previous studies reported the *Eola1*’s cardiac-enriched expression^24^, prompting us to investigate its impact on mitochondrial function in cardiomyocytes. Strikingly, while steady-state mt-mRNA levels remained unchanged (Extended Data Fig. 4e), *Eola1*^-/-^ mice exhibited a pronounced decrease in ATP6, ND5, and COX1 protein levels compared to *Eola1*^+/+^ mice in cardiomyocytes (Fig. 4b). To thoroughly characterize the cardiac consequences of *Eola1* deficiency, we performed in-depth functional assessments of hearts through transthoracic echocardiography in young adult mice. Results demonstrated that compared to *Eola1*^+/+^ mice, *Eola1*^-/-^ mice exhibited cardiac enlargement with diminished systolic function, phenocopying the clinical manifestations of dilated cardiomyopathy (Fig. 4c-l). In summary, our study demonstrates that EOLA1 is a key regulator of mitochondrial translation through its interaction with 12S mt-rRNA and TUFM, thereby sustaining OXPHOS-dependent bioenergetics and maintaining normal cardiac function (Fig. 5).

**Fig. 4:**
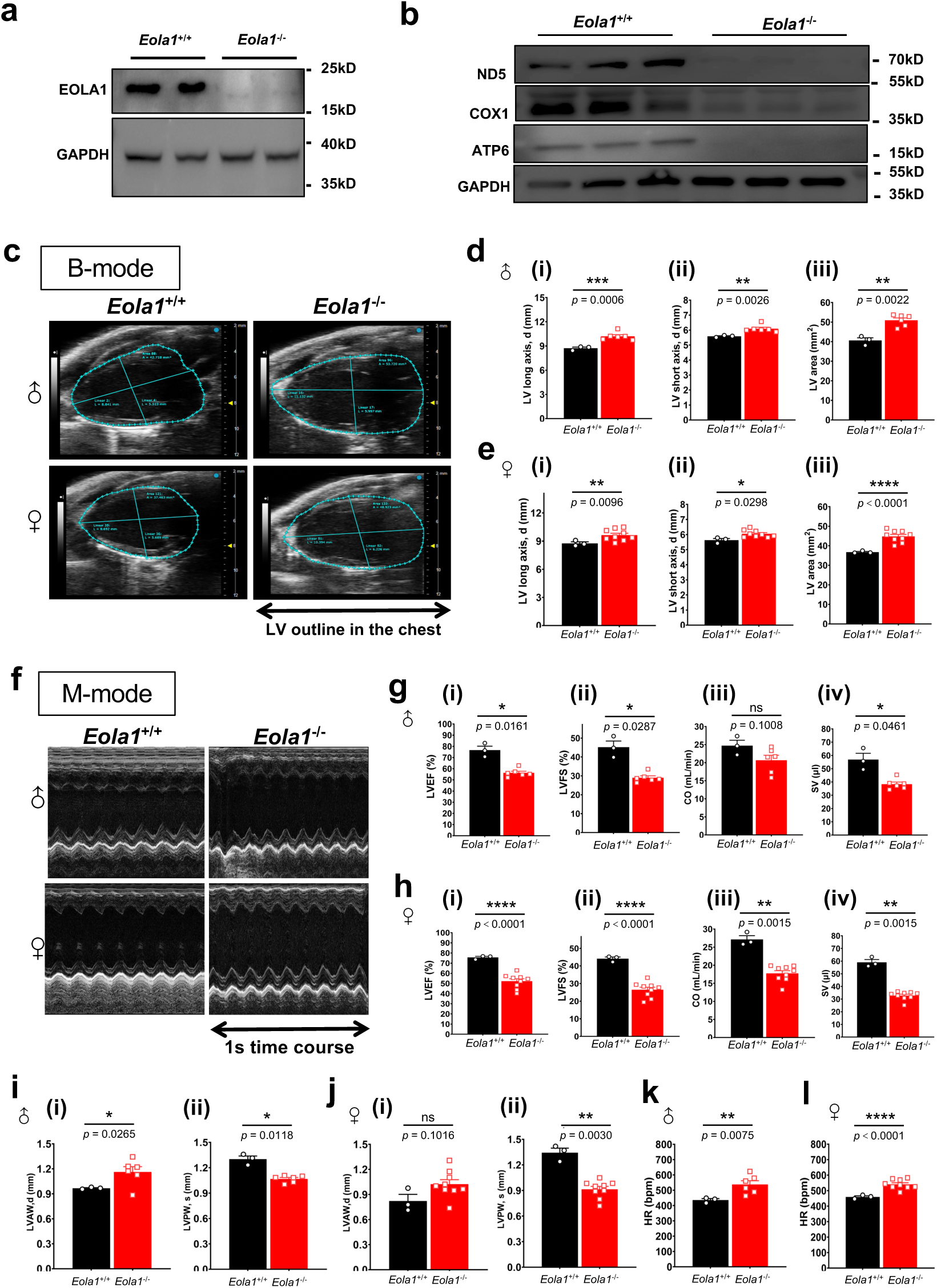
Abnormal cardiac function was observed in *Eola1*^-/-^ mice. **a,** EOLA1 protein deficiency in knockout mice was validated by Western blotting analysis. GAPDH was used as a loading control. **b,** Western blotting analysis revealed significantly lower levels of mitochondrial protein in *Eola1*^-/-^ hearts compared to *Eola1***^+/+^**. GAPDH was used as a loading control. **c,** Typical ventricular borders at end-diastole were identified by B-mode echocardiography. **d, e,** Compared to *Eola1***^+/+^** mice, both male (**d**) and female (**e**) *Eola1*^-/-^ mice demonstrated significant enlargement in ventricular long-axis length (**i**), short-axis length (**ii**), and cross-sectional area (**iii**). **f,** Representative images of left ventricular motion patterns identified by M-mode echocardiography. **g,** Compared to *Eola1***^+/+^** mice, male *Eola1*^-/-^ mice demonstrated a significant decrease in ejection fraction (**i**), shortening fraction (**ii**), and stroke volume (**iv**) of left ventricle. Cardiac output (**iii**) of left ventricle also showed a decreasing trend. **h,** Compared to *Eola1***^+/+^** mice, female *Eola1*^-/-^ mice demonstrated a significant decrease in ejection fraction (**i**), shortening fraction (**ii**), cardiac output (**iii**) and stroke volume (**iv**) of left ventricle. **i,** *Eola1* knockout leads to thickening of the anterior wall (**i**) and thinning of the posterior wall (**ii**) of the left ventricle in male mice. **j,** *Eola1* deficiency causes thinning of the posterior wall (**ii**) and a trend of thickening of the anterior wall (**i**) of the left ventricle in female mice. **k, l,** *Eola1* deficiency increases heart rate of both male (**k**) and female (**l**) mice. **Note: d**, **e**, **g-l**, two-tailed unpaired Student’s t-tests. \**P*<0.05, \*\**P*<0.01, \*\*\**P*<0.001, \*\*\*\**P*<0.0001, ns: not significant. Data were presented as mean ± SEM.

**Fig. 5:**
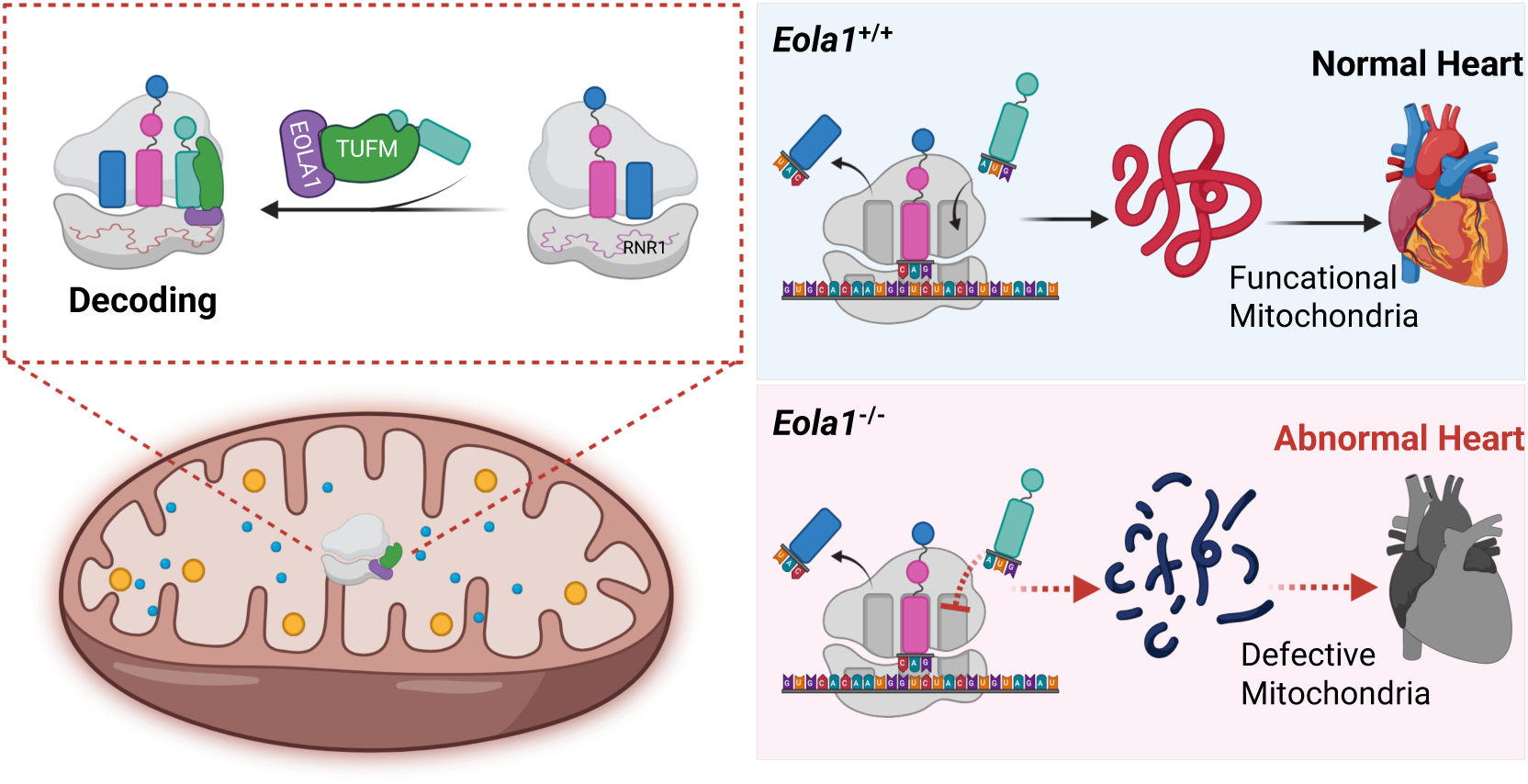
Working model illustrating the critical role of mitochondrial protein EOLA1 in heart function. EOLA1, a mitochondrial matrix protein identified by CRISPR screening, interacts with TUFM/12S mt-rRNA to promote the protein synthesis of OXPHOS subunits. Its loss impairs mt-mRNA translation and causes heart failure in mice, revealing a mitochondrial translation-cardiac function axis pivotal for cardiovascular homeostasis.

## DISCUSSION

In this study, we identified EOLA1 as an essential regulator of mitochondrial metabolic homeostasis through an unbiased functional screening. Our integrative analysis combining bioinformatics and experimental validation reveals EOLA1 as a previously uncharacterized mitochondrial matrix protein that physically binds both 12S mt-rRNA and elongation factor TUFM, thereby enhancing the translation elongation efficiency of mitochondrial-encoded mRNAs. Genetic ablation of *Eola1* in mammalian cells leads to a marked impairment of mitochondrial protein synthesis and a consequent reduction in OXPHOS capacity, highlighting its essential role in maintaining mitochondrial bioenergetics. Importantly, *Eola1* knockout mice display pronounced cardiac dysfunction, supporting a critical role for EOLA1 in sustaining cardiac homeostasis via regulation of mitochondrial translation. Together, these findings establish EOLA1 as a key modulator of mitochondrial gene expression and reveal a previously unappreciated translational mechanism underlying mitochondrial dysfunction in cardiovascular disease.

### Structural and functional insights into the ASCH domain of EOLA1

EOLA1 was initially identified as a LPS responsive gene in endothelial cells^49^, and was later shown to regulate inflammatory responses through its physical interaction with MT2A^24^. This interaction stabilizes MT2A protein levels, resulting in the suppression of IL-6^25^ and VCAM-1^26^ expression, and attenuation of downstream LPS signaling pathways. This molecular mechanism may contribute to inflammatory diseases such as diabetic foot ulcers^50^. Structurally, EOLA1 is an ASCH domain-containing protein with a characteristic β-barrel structure flanked by α-helices, containing a conserved “GxKxxExR” motif in its substrate-binding cavity^22^. The ASCH domain, a member of the PUA-like superfamily, plays versatile roles in RNA metabolism during transcriptional coactivation, RNA processing, and translation regulation. While a hypothetical ASCH-containing protein from *Zymomonas mobilis* functions as a monomeric ribonuclease that binds and degrades single-stranded RNA *in vitro*^28,29^, *E. coli* EcYqfB exhibits amidohydrolase activity toward ac^4^C modification, while lacking detectable RNA-binding or ribonuclease functions^51,52^. Humans encode three ASCH domain-containing proteins, TRIP4, EOLA1 (CXORF40A), and EOLA1 paralog EOLA2 (CXORF40B)^23^. Among them, TRIP4 (also known as ASC1) was initially implicated in regulating transcription as part of the tetrameric nuclear activating signal cointegrator complex, which is a putative RNA-interacting complex^53,54^. Structural studies further reveal that TRIP4’s ASCH domain engages double-stranded DNA via two adjacent positively charged surface patches in a sequence-independent manner^55^. However, more recent studies demonstrated that TRIP4 mediated subunit dissociation of collided ribosome during RQC (Ribosome-associated Quality Control)^56–58^. Consistent with these latest findings, our current study also indicated that EOLA1 specifically binds 12S mt-rRNA and modulates mitochondrial translation, implying a likely conserved role of ASCH domain in translation regulation

### EOLA1 as a novel mitochondrial translation regulator

Mitochondria evolved from proteobacteria and inherited bacterial-like translational machinery. However, their protein synthesis system has evolved unique features through adaptation, including specialized mitoribosomes, distinct translation factors, modified tRNAs and mRNAs, and altered genetic codon usage^6,59^. The mitochondrial translation process consists of four key steps: initiation, elongation, termination, and recycling. Among these, elongation is the most evolutionarily conserved phase, where elongation factors exhibit homology across bacteria, eukaryotic cytosol, and mitochondria^60,61^. For example, mitochondrial elongation factor TUFM (also known as mtEF-Tu), is a highly conserved GTPase that shares significant structural homology with bacterial EF-Tu^62^. It delivers aminoacyl-tRNA to the ribosomal A-site in its GTP-bound form, an energy-dependent process coupled with GTP hydrolysis. Subsequently, the inactive mtEF-Tu-GDP complex is reactivated to mtEF-Tu-GTP through the action of the guanine nucleotide exchange factor (mtEF-Ts), enabling it to re-enter the tRNA delivery cycle^6,60,61^. In our study, we identified EOLA1 as a mitochondrial matrix protein that directly binds 12S mt-rRNA and further demonstrated that EOLA1 modulates mitochondrial translation through its interaction with TUFM. Functional studies reveale<u>d</u> that EOLA1 deficiency impairs mitochondrial translation, resulting in diminished synthesis of OXPHOS subunits and attenuated respiratory chain activity. In future work, structural characterization of the EOLA1/mtEF-Tu/mitoribosome interaction is required to elucidate the precise binding interface and regulatory role of EOLA1 in mitochondrial translation.

### Heart disease and dysregulation of mitochondrial translation

The adult human heart produces and consumes ∼1 kg of ATP daily to sustain its contractile function, with mitochondrial OXPHOS accounting for over 95% of ATP generation^6,16,17,63^. This immense energy demand is met through a highly specialized mitochondrial network, which occupies over 40% of cardiomyocyte volume, densely packed between myofibrils, around the nucleus, and beneath the sarcolemma^64^. Given this critical reliance on mitochondria, dysfunction in these organelles is a major contributor to cardiac disease. Indeed, Mutations in genes regulating mitochondrial translation including those in mt-DNA as well as nuclear genes encoding ribosomal proteins, translation factors, or tRNA-modifying enzymes, are associated with cardiomyopathy in over 50% of all carriers^18^. Our current study demonstrates that *EOLA1* deficiency disrupts mitochondrial translation, leading to severe OXPHOS dysfunction, impaired ATP production, and ultimately metabolic dysfunction in cardiomyocytes. Notably, *EOLA1*-knockout mice recapitulate these defects, underscoring the non-redundant role of mitochondrial translation in cardiac homeostasis. These findings establish mitochondrial protein synthesis as a fundamental vulnerability in heart disease and highlight potential therapeutic strategies targeting this pathway.

In summary, we demonstrate that EOLA1, a novel mitochondrial matrix protein, enhances translation elongation by binding 12S mt-rRNA and TUFM to maintain metabolic homeostasis. Loss of EOLA1 disrupts mitochondrial protein synthesis, impairing OXPHOS in cells and triggering cardiac dysfunction in mice. This identifies EOLA1 as a key translational regulator linking mitochondrial gene expression to cardiovascular health.

## ACKNOWLEDGEMENTS

We would like to thank all members in Hao Chen’s Lab for their help and advice in experimental design. We thank Dr. Ruilin Tian, Dr. Honghui Ma, Dr. Pang-Hung Hsu, Dr. Marcia Haigis and Dr. Alison E. Ringel for all the kind suggestions and technical assistance. The experiments presented in Figures 2b and 3a were conducted at Harvard Medical School, with which Yang Shi and Hao Chen were previously affiliated.

## AUTHOR CONTRIBUTIONS

Y.S. and H.C. designed and conceived the experiments. X.Y.S. and Y.Y.Z. performed most experiments. R.Q.W. and N.Z. performed complex purification and RIP-qPCR. N.B.L. and S.J.Z performed the mouse experiments under the supervision of P.Z.. H.C., R.Q.W. and Y.L.C. assisted with polysome profiling. D.W. and Y.X.J. assisted with cell culture. S.M.F. and J.Y. helped to prepare figures. H.C. and C.H.H analyzed the data. X.Y.S., H.C. and Y.S. wrote the manuscript. All authors have read and approved the final manuscript.

## DATA AND MATERIALS AVAILABILITY

The raw data of NGS is available in the GEO database under the accession number: GSE300159 (token: sleruykotxylvab). The mass spectrometry proteomics data have been deposited to the ProteomeXchange Consortium via the iProX partner repository with the dataset identifier PXD066586. The proteome datasets can be accessed at https://www.iprox.cn/page/PSV023.html;?url=1759505133475Trez using password kdnW.

---

**Extended Data Fig. 1:**
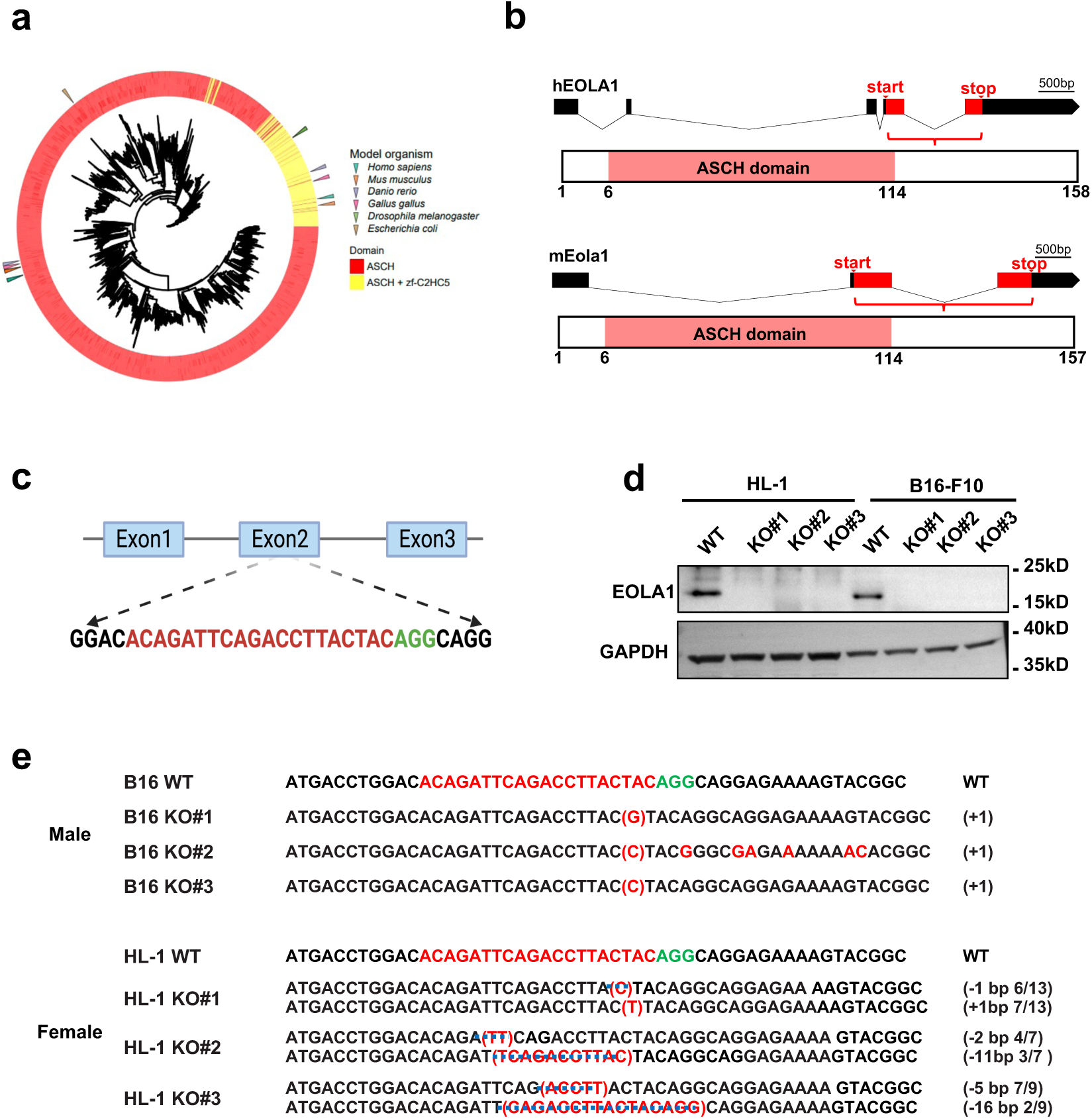
Generation and validation of *Eola1*-knockout cell lines. **a,** Evolutionary analysis of the ASCH domain reveals high conservation across species. **b,** A scheme showing functional domains of EOLA1 in *Homo sapiens* (top) and *Mus musculus* (bottom). The ASCH domain is shown in red. **c,** Schematic diagram of sgRNA targeting mouse *Eola1* locus. **d,** Western blotting analysis confirming *Eola1* knockout clones in B16-F10 and HL-1 cell lines with anti-EOLA1 antibody. GAPDH was used as a loading control. **e,** Sanger sequencing results of *Eola1* KO cell lines with genetic mutations introduced with the CRISPR/Cas9 system.

**Extended Data Fig. 2:**
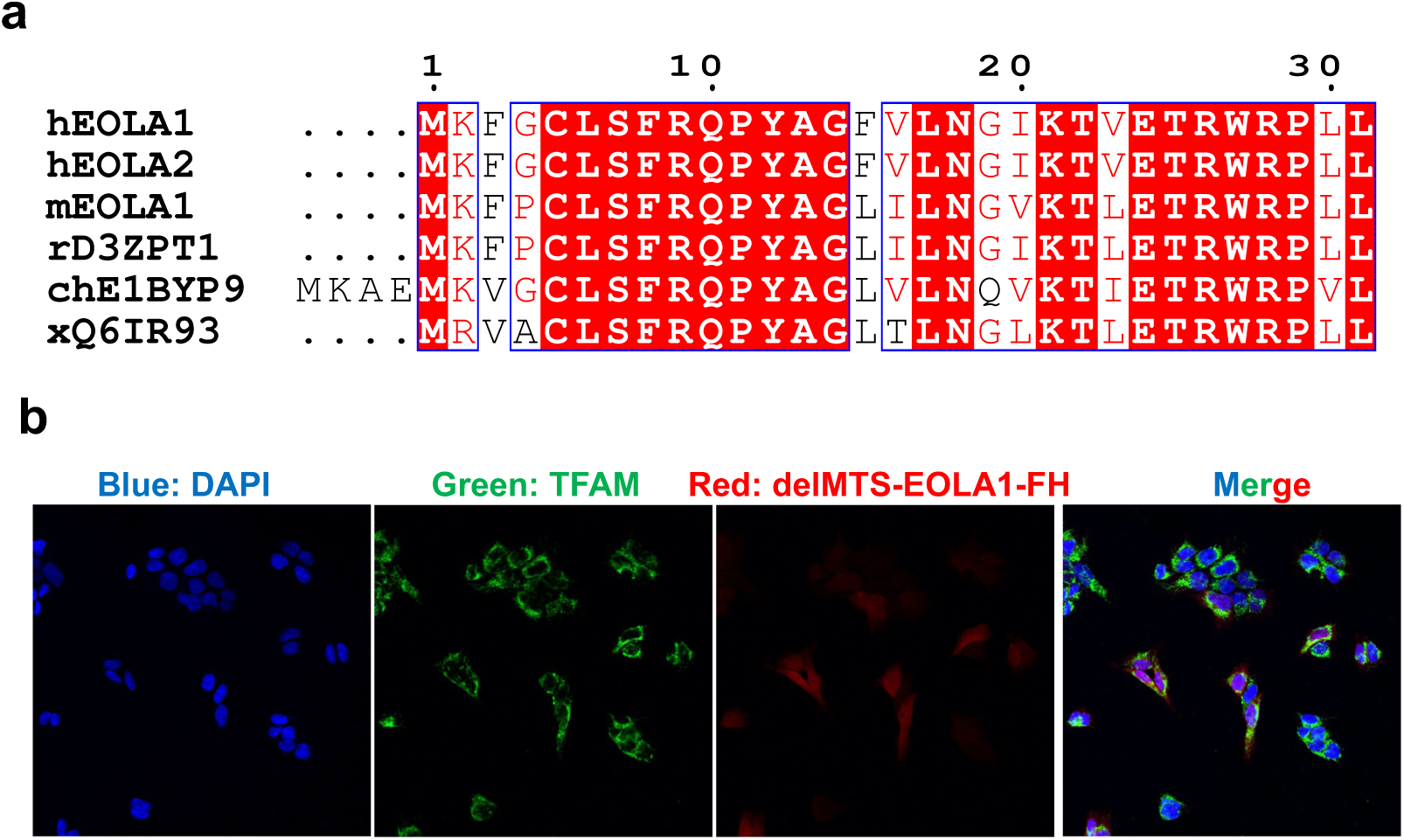
Mislocalization of EOLA1 upon deletion of its N-terminal targeting signal. **a,** Evolutionary conservation of EOLA1’s N-terminal domain. Sequence alignment across diverse species reveals key conserved residues (red). **b,** Fluorescence imaging analysis of EOLA1 subcellular localization after N-terminal mitochondrial targeting signal (MTS) deletion (red). Mitochondria were labeled with TFAM (green). Exogenously expressed MTS-deficient EOLA1 (delMTS-EOLA1) carried a C-terminal FLAG-HA tag.

**Extended Data Fig. 3:**
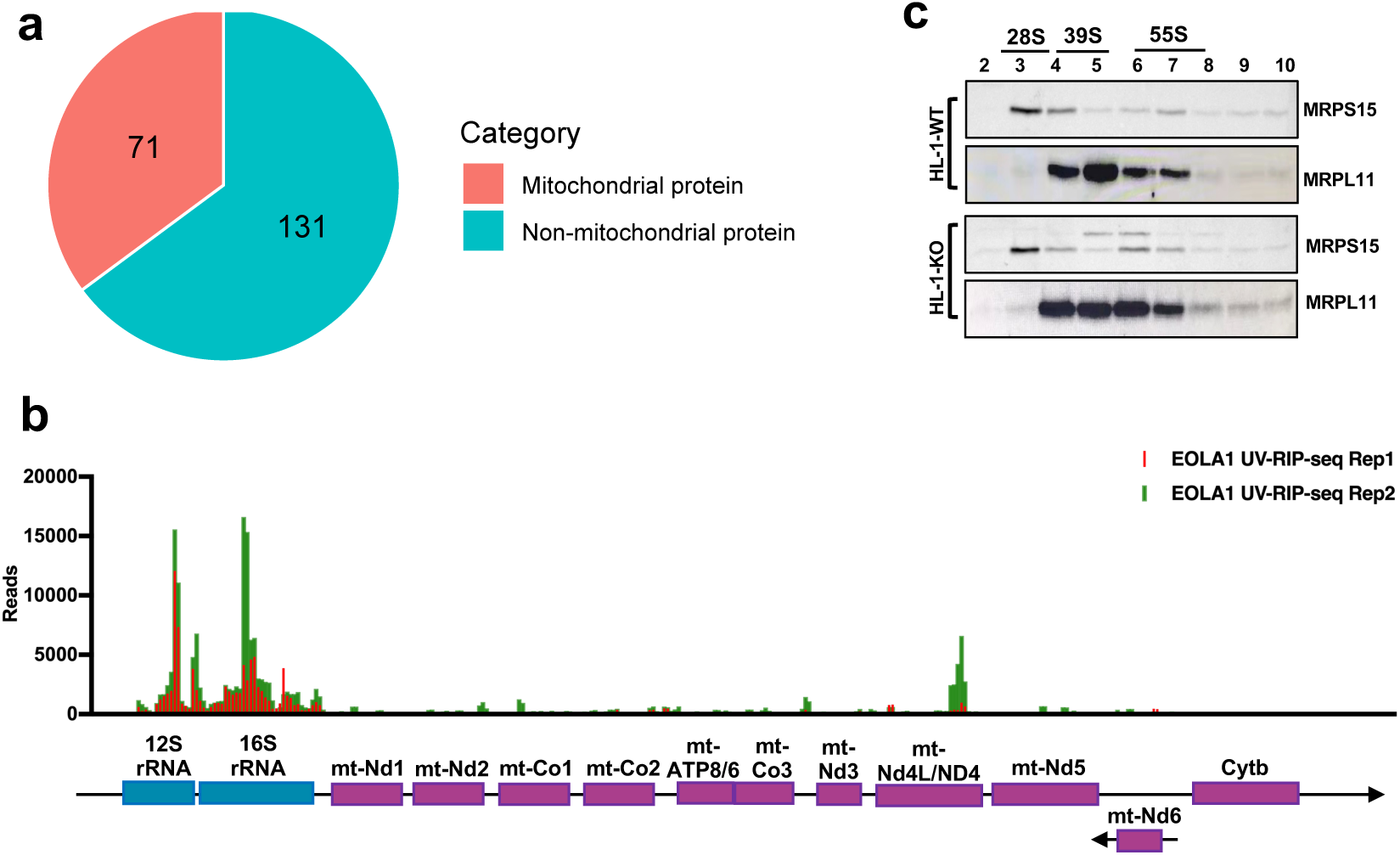
EOLA1 is associated with mitochondrial translation. **a,** Pie chart representing the proportion of mitochondrial proteins identified in the EOLA1 interactome. **b,** EOLA1-interacting mitochondrial RNAs were identified via UV-RIP-seq. **c,** Protein levels of MRPS15 (mt-SSU marker) and MRPL11 (mt-LSU marker) were analyzed by Western blotting across fractions.

**Extended Data Fig. 4:**
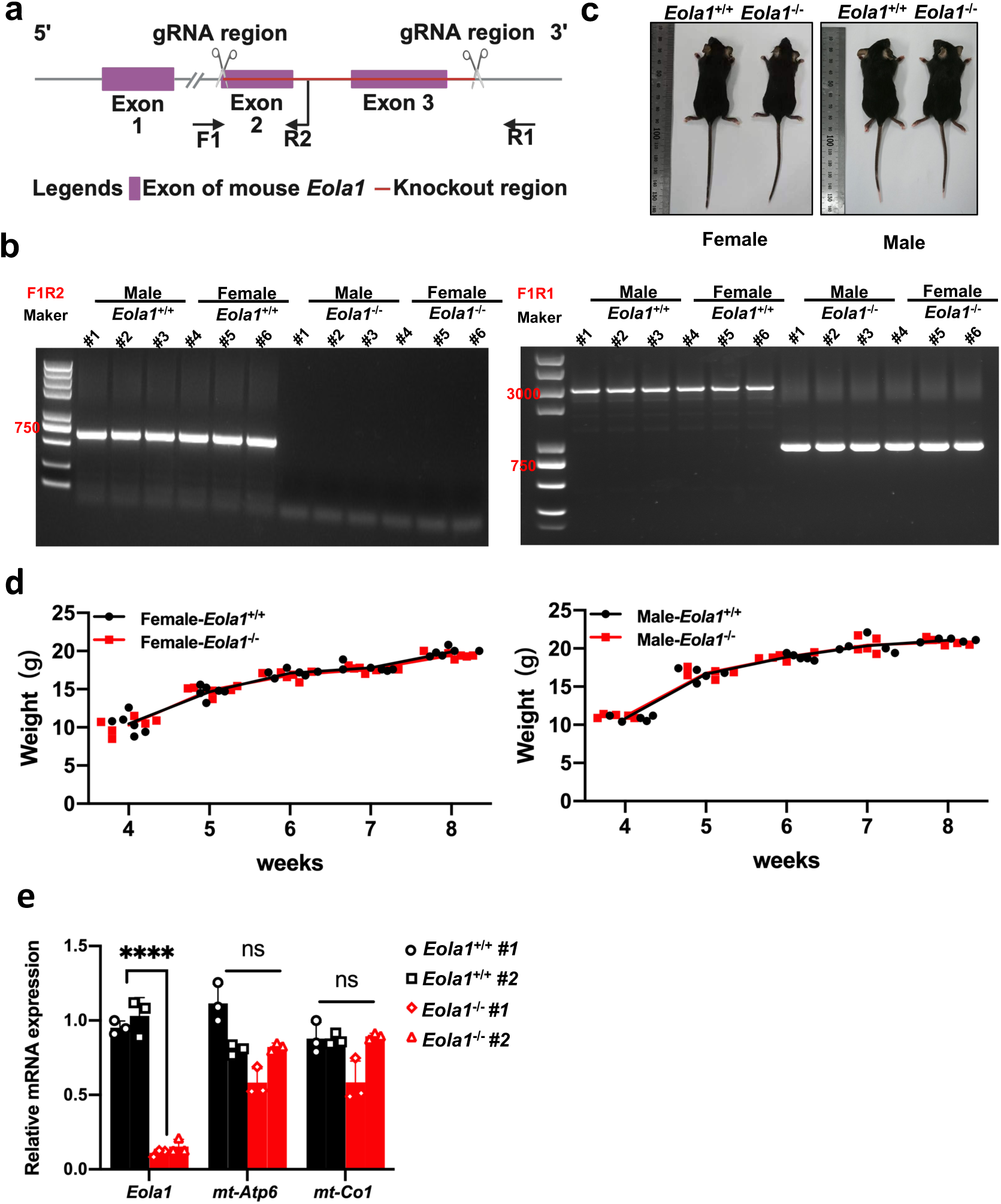
Generation and phenotypic analysis of *Eola1*^-/-^ mice. **a, b,** Schematic diagram of sgRNA targeting mouse *Eola1* locus (**a**) and validation of *Eola1* knockout in mice by PCR (**b**). The genotyping primers are provided in supplementary Table 1. **c,** Representative pictures of the *Eola1*^+/+^ and *Eola1*^-/-^ mice. **d,** Comparison of body weight in female (left) and male (right) *Eola1*^+/+^ and *Eola1*^-/-^ mice. **e,** Relative mRNA levels of mitochondrial-encoded transcripts were quantified by RT-qPCR in *Eola1*^+/+^ and *Eola1*^-/-^ hearts. **Note: e**, two-tailed unpaired Student’s t-tests. \*\*\*\**P*<0.0001, ns: not significant. Data were presented as mean ± SD (n=6).

**Supplementary Table S1.**
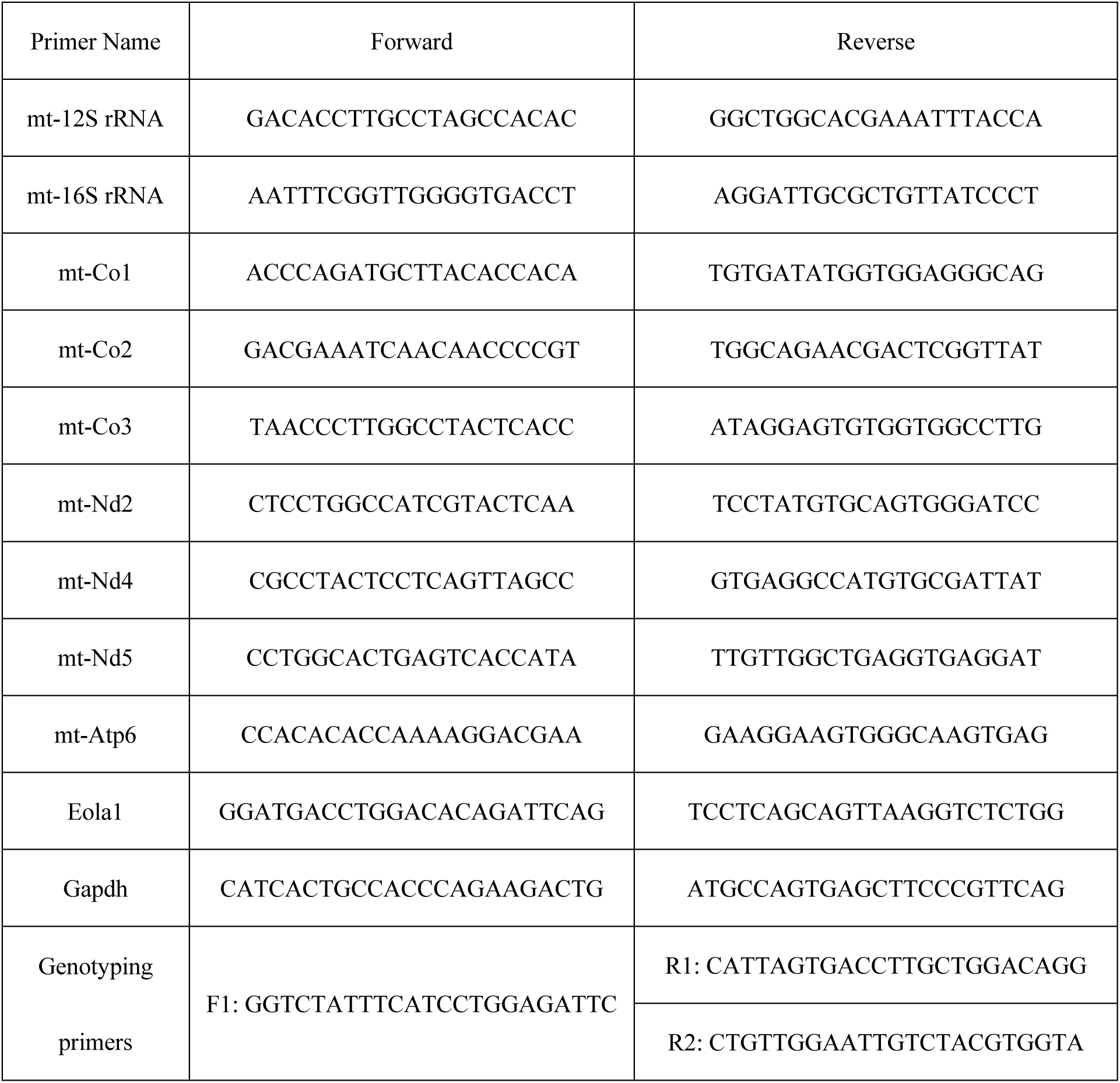
List of PCR primers used in this study.

